# Evasion of Neutralizing Antibody Response by the SARS-CoV-2 BA.2.75 Variant

**DOI:** 10.1101/2022.08.14.503921

**Authors:** Panke Qu, John P. Evans, Yi-Min Zheng, Claire Carlin, Linda J. Saif, Eugene M. Oltz, Kai Xu, Richard J. Gumina, Shan-Lu Liu

## Abstract

The newly emerged BA.2.75 SARS-CoV-2 variant exhibits an alarming 9 additional mutations in its spike (S) protein compared to the ancestral BA.2 variant. Here we examine the neutralizing antibody escape of BA.2.75 in mRNA-vaccinated and BA.1-infected individuals, as well as the molecular basis underlying functional changes in the S protein. Notably, BA.2.75 exhibits enhanced neutralization resistance over BA.2, but less than the BA.4/5 variant. The G446S and N460K mutations of BA.2.75 are primarily responsible for its enhanced resistance to neutralizing antibodies. The R493Q mutation, a reversion to the prototype sequence, reduces BA.2.75 neutralization resistance. The mutational impact is consistent with their locations in common neutralizing antibody epitopes. Further, the BA.2.75 variant shows enhanced cell-cell fusion over BA.2, driven largely by the N460K mutation, which enhances S processing. Structural modeling revealed a new receptor contact introduced by N460K, supporting a mechanism of potentiated receptor utilization and syncytia formation.

## Introduction

Emergence of the Omicron variant of SARS coronavirus 2 (SARS-CoV-2) in late 2021 sparked an unprecedented wave of coronavirus disease 2019 (COVID-19) cases and exhibited robust evasion of vaccine- and infection-induced immunity (Gruell et al., 2022; Hoffmann et al., 2022). More recently, several subvariants of Omicron have been identified, which have driven subsequent waves of infection. The BA.1 subvariant, responsible for the initial Omicron wave, was replaced by BA.2, which displayed slightly enhanced transmissibility and resistance to BA.1-induced sera (Centers for Disease Control and Prevention, 2022; Evans et al., 2022; Yamasoba et al., 2022b). BA.2 then evolved into several progeny subvariants, including the BA.2.12.1 variant, which subsequently became predominant (Centers for Disease Control and Prevention, 2022). Remarkably, the BA.4 and BA.5 variants, which bear identical spike (S) proteins and evolved from BA.2, are currently dominant in the world, including in the US (Centers for Disease Control and Prevention, 2022). BA.4 and BA.5 bear an L452R mutation that is primarily responsible for further enhanced neutralizing antibody resistance (Qu et al., 2022; Tuekprakhon et al., 2022). Recently, another distinct BA.2-derived subvariant, BA.2.75, has been identified. BA.2.75 is increasing in prevalence in southeast Asia and has been detected globally (Callaway, 2022). Notably, BA.2.75 bears 9 key S mutations including K147E, W152R, F157L, I210V, G257S, D339H, G446S, and N460K, as well as an R493Q reversion mutation (World Health Organization, 2022) (**Fig. 1A**). These mutations, particularly those in the receptor binding domain (RBD), have generated concern over further immune escape.

**Figure 1:**
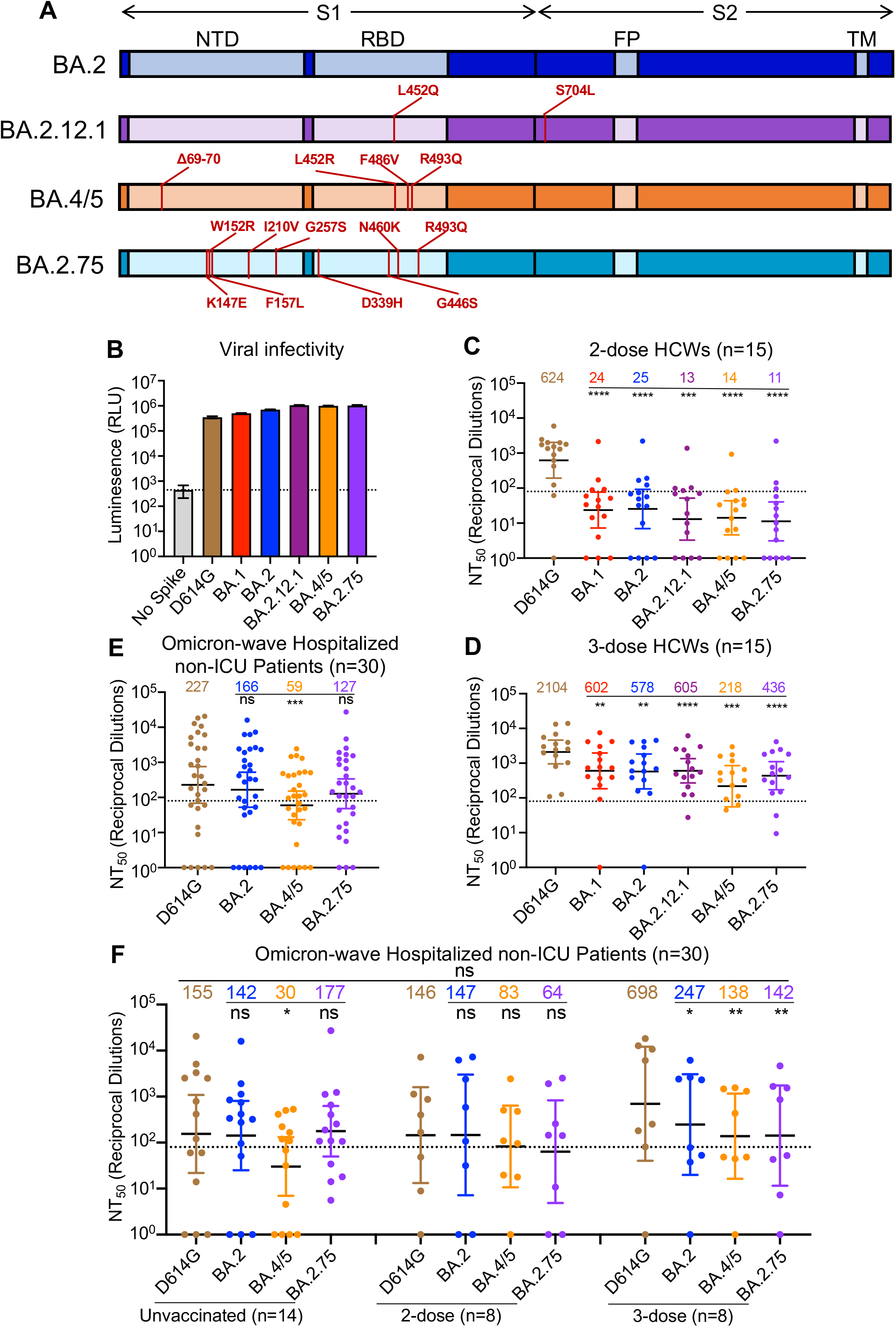
BA.2.75 exhibits strong neutralization resistance to 2-dose and 3-dose mRNA vaccinee sera and Omicron wave patient sera. (**A**) Schematic of BA.2-derived SARS-CoV-2 variants with mutations relative to the BA.2 background indicated. Highlighted are the S1 and S2 subunits, N-terminal domain (NTD), receptor binding domain (RBD), fusion peptide (FP), and transmembrane domain (TM). (**B**) Infectivity of pseudotyped lentivirus bearing S protein from SARS-CoV-2 variants of study; bars represent means ± standard error. (**C-D**) Neutralizing antibody titers against lentivirus pseudotyped with S from individual SARS-CoV-2 variants for 15 health care workers for sera collected 3-4 weeks after second mRNA vaccination (C) or 1-12 weeks after homologous mRNA booster vaccination (D). (**E**) Neutralizing antibody titers for sera collected from 30 COVID-19 patients hospitalized during the BA.1 pandemic wave. (**F**) Neutralizing antibody titers against hospitalized BA.1 wave patients are divided by vaccination status. (C-F) Dots indicate individual patient samples; bars represent geometric means with 95% confidence intervals; significance relative to D614G was determined by one-way repeated measures ANOVA with Bonferroni multiplicity correction. P-values are displayed as *p < 0.05, **p < 0.01, ***p < 0.001, ****p < 0.0001, and ns for not significant.

Here we characterize the BA.2.75 S protein by examining its sensitivity to neutralizing antibodies from mRNA-vaccinated and/or boosted health care workers (HCWs), as well as from Omicron-wave-hospitalized COVID-19 patients. In addition, we examine BA.2.75 infectivity, S processing, and fusogenicity. Mutational analysis revealed the N460K as a key driver of enhanced fusogenicity, while the G446S and N460K mutations were primarily responsible for reduced neutralization sensitivity of BA.2.75 compared to BA.2. Moreover, we find that the R493Q reversion mutation enhances the neutralization sensitivity of BA.2.75. These findings inform our understanding of SARS-CoV-2 evolution and will aid in addressing the ongoing threat of emerging SARS-CoV-2 variants.

## Results

### BA.2.75 exhibits enhanced neutralization resistance over BA.2

We first sought to characterize sensitivity to vaccine-induced immunity of the BA.2.75 variant. Utilizing our previously reported pseudotyped lentivirus assay (Zeng et al., 2020), we examined neutralizing antibody (nAb) titers for 15 Ohio State University Wexner Medical Center health care workers (HCWs) in serum samples collected 3-4 weeks after vaccination with a second dose of Moderna mRNA-1273 (n = 7) or Pfizer/BioNTech BNT162b2 (n = 8) vaccine, and 1-12 weeks after vaccination with a homologous booster dose (see STAR Methods). Patient sera were examined for nAb titers against lentivirus pseudotyped with S from ancestral SARS-CoV-2 S bearing only the D614G mutation, as well as S from BA.1, BA.2, BA.2.12.1, BA.4/5, and BA.2.75 (Fig. 1A). All S constructs were functional and produced comparably infectious lentivirus pseudotypes (Fig. 1B).

Notably, all Omicron sublineages, including BA.2.75, exhibited strong resistance to 2-dose-induced immunity compared to D614G (p < 0.0001), with only 1-2 HCW samples exhibiting 50% neutralization titers (NT_50_) above the limit of quantification (NT_50_ = 80) (Fig. 1C). In contrast, administration of a booster dose recovered the neutralizing antibody response against all Omicron subvariants (Fig. 1D, Fig. S1A-H). In serum from the boosted individuals, BA.2.75 exhibited 4.8-fold (p < 0.0001) lower neutralization than D614G, with somewhat stronger neutralization resistance than BA.2 and BA.2.12.1, which were neutralized 3.6-fold (p < 0.01) and 3.5-fold (p < 0.001) less efficiently than D614G, respectively (Fig. 1D). However, BA.2.75 showed higher neutralization sensitivity than BA.4/5, which had 9.7-fold (p < 0.001) lower neutralization than D614G (Fig. 1D).

We also examined the neutralizing antibody response in a cohort of non-ICU COVID-19 patients (n = 30) hospitalized at the Ohio State University Wexner Medical Center during the Omicron-wave of the pandemic. These patient samples were collected between early February and early March of 2022, representing a BA.1 dominant period in Ohio. Overall, the nAb titers of the Omicron-wave patients were much lower than those of boosted HCWs, and BA.2.75 exhibited neutralization resistance modestly higher than BA.2 (by 44.0%, p > 0.05) but much lower than BA.4/5 (3.8-fold; p < 0.001) relative to D614G (Fig. 1E; Fig. S1I). This cohort of Omicron-wave patients included 14 unvaccinated patients, 8 patients vaccinated with 2 doses of Moderna mRNA-1273 (n = 4) or Pfizer/BioNTech BNT162b2 (n = 4), and 8 patients vaccinated and boosted with Pfizer/BioNTech BNT162b2. We found that, while BA.2.75 was neutralized comparably to BA.2 and D614G for unvaccinated patients, BA.2.75 was neutralized 2.3-fold less efficiently than D614G in 2-dose vaccinated patients (p > 0.05) and 4.9-fold less efficiently for 3-dose vaccinated patients (p < 0.01), respectively (Fig. 1F). The boosted HCWs with breakthrough infection exhibited higher nAb titers overall (Fig. 1F), as would be expected.

### BA.2.75 neutralization is modulated by G446S, N460K, and R493Q mutations

To understand the determinants of BA.2.75 neutralization resistance, we examined all nine point mutations in the BA.2 background, as well as nine corresponding reversion mutations in the background of BA.2.75. None of these single mutations substantially impacted lentiviral pseudotype infectivity (Fig. 2A-B). We then examined the neutralization sensitivity of these mutants to sera from 9 HCWs collected 1-12 weeks after homologous booster vaccination with Moderna mRNA-1273 (n = 2) or Pfizer/BioNTech BNT162b2 (n = 7). When the G446S mutation was introduced to BA.2, a slight but significant reduction in sensitivity to 3-dose mRNA vaccine-induced nAbs was observed (42.7%, p < 0.01), which was comparable to BA.2.75 (Fig. 2C). Introduction of a S446G reversion mutation into BA.2.75 enhanced neutralization sensitivity by 31.4%, albeit the change was not statistically significant (p = 0.055) (Fig. 2D). Interestingly, introduction of a R493Q mutation into BA.2 increased neutralization sensitivity by 35.8% (p > 0.05), while introduction of the Q493R reversion mutation into BA.2.75 reduced neutralization sensitivity by 45.1% (p > 0.05) (Fig. 2C-D). Of note, the N460K mutation also substantially increased neutralization resistance of BA.2 by 33.0% (p > 0.05), whereas the K460N reversion mutation in BA.2.75 was 77.4% (p = 0.069) more neutralization sensitive (Fig. 2C-D). Thus, the G446S and N460K mutations in BA.2.75 are largely responsible for its enhanced neutralization resistance, while the R493Q reversion mutation in BA.2.75 at least partially restores neutralizing epitopes found in the prototype SARS-CoV-2, which were otherwise abolished in BA.2.

**Figure 2:**
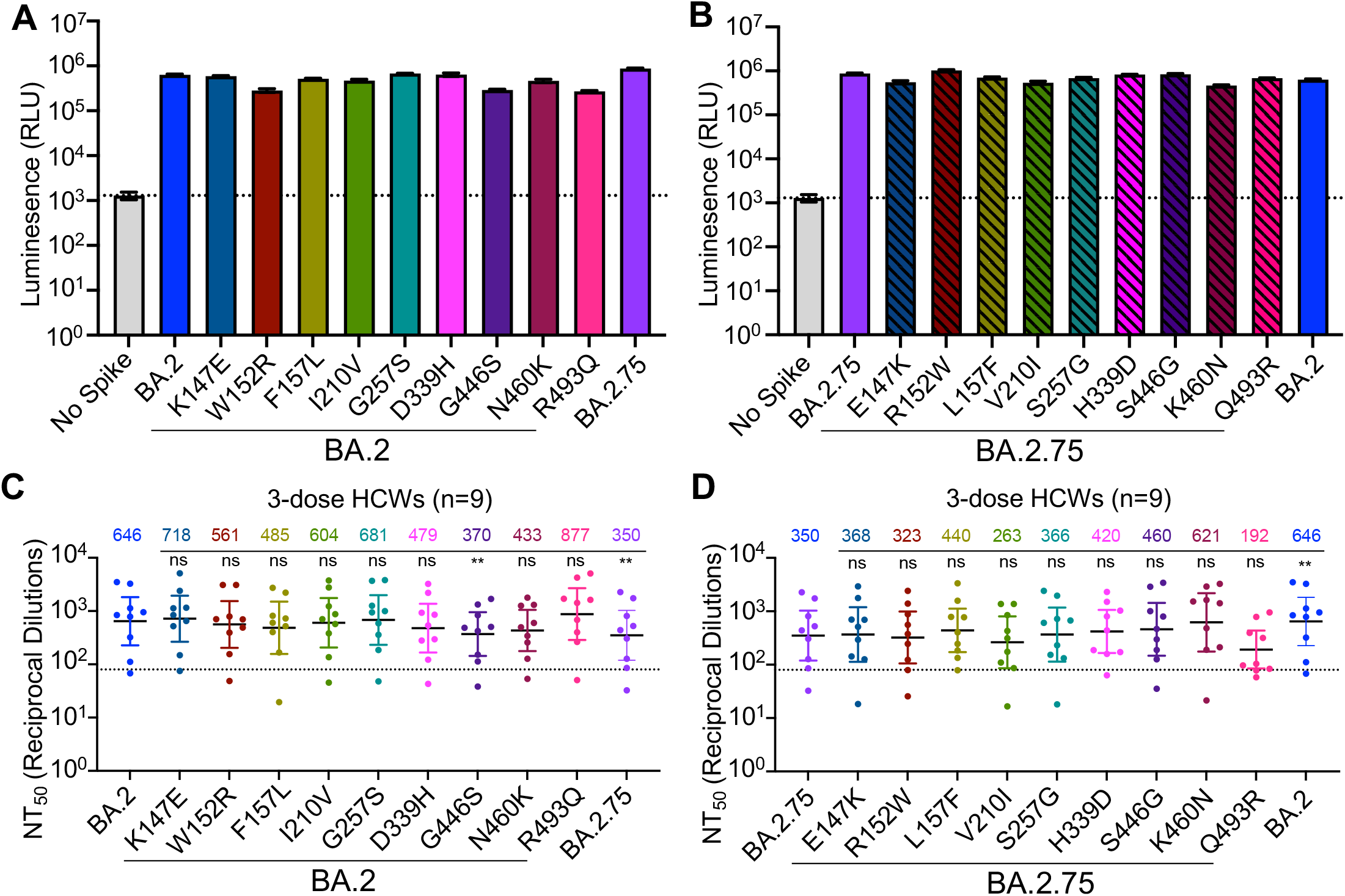
The G446S, N460K, and R493Q mutations modulate BA.2.75 neutralization sensitivity. (**A**) Relative infectivity of lentivirus pseudotyped with BA.2 S with single mutations from BA.2.75 lineage defining mutations; bars represent means ± standard error. (**B**) Relative infectivity of lentivirus pseudotyped with BA.2.75 S with single reversion mutations to remove BA.2.75 lineage defining mutations; bars represent means ± standard error. (**C-D**) Neutralizing antibody titers against lentivirus pseudotyped with S from BA.2 with single mutations from BA.2.75 lineage-defining mutations (C) or BA.2.75 with single reversion mutations from BA.2.75 lineage-defining mutations (D) for sera collected from 9 health care workers 1-12 weeks after homologous mRNA booster vaccination. Dots indicate individual patient samples; bars represent geometric means with 95% confidence intervals; significance relative to D614G was determined by one-way repeated measures ANOVA with Bonferroni multiplicity correction. P-values are displayed as **p < 0.01, and ns for not significant.

### BA.2.75 exhibits enhanced syncytia formation and S processing compared to BA.2

We next sought to characterize key features of the BA.2.75 S protein, including the ability to mediate cell-cell fusion. HEK293T-ACE2 cells were transfected to express GFP and variant SARS-CoV-2 S proteins. As previously reported (Zeng et al., 2021), all Omicron sublineages exhibited reduced fusogenicity compared to the ancestral D614G S (Fig. 3A-B). However, BA.2.75 exhibited enhanced syncytia formation compared to BA.2, with mean syncytia size 2.0-fold higher than BA.2 (p < 0.0001) (Fig. 3A-B; Fig. S2A); this was despite similar surface expression, as examined by flow cytometry (Fig. 3C-D).

**Figure 3:**
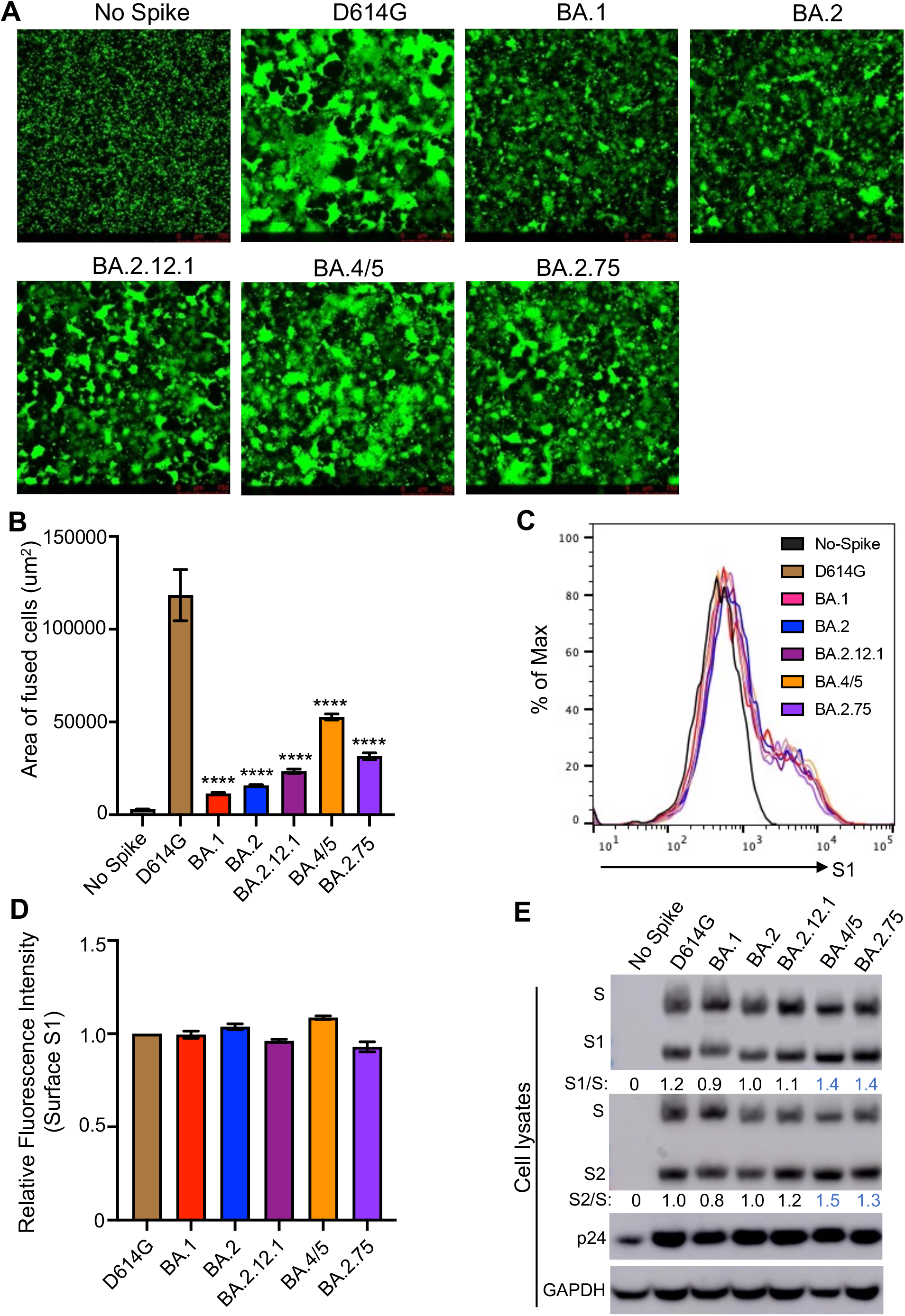
BA.2.75 exhibits enhanced cell-cell fusion and S processing. (**A**) Fluorescence images displaying syncytia formation are presented for HEK293T-ACE2 cells 48 hr after co-transfection with a GFP expression construct and SARS-CoV-2 variant S proteins. (**B**) Quantification of syncytia formation in panel (A) displays the mean syncytia size; bars represent means ± standard error, with significance relative to D614G determined by one-way ANOVA with Bonferroni multiplicity correction. (**C**) Histogram displays of the surface staining of HEK293T cells expressing S proteins, which were detected by an anti-S1 antibody (T62). (**D**) Quantification of relative surface expression as shown in (C); bars represent means ± standard error. (**E**) Pseudotyped lentivirus producer cell lysate was assessed for processing of S by probing with anti-S1 (T62), anti-S2, anti-HIV-1 Gag (anti-p24), and anti-GAPDH. Band intensities were quantified in ImageJ and the ratio of S1/S or S2/S is displayed relative to the S1/S or S2/S ratio of BA.2. P-values are displayed as ****p < 0.0001.

To determine if the enhanced fusogencity phenotype might be related to alterations in processing of S protein, we examined cell lysates from the pseudotyped lentivirus producer. As shown in Figure 3E, BA.2.75 spike exhibited enhanced processing, as reflected in the ratio of S1 or S2 subunit to full length S ratio, which was ∼30-40% higher than BA.2. Consistent with its enhanced fusion, BA.4/5 showed the highest S processing among omicron variants (Fig. 3A-B and E; Fig. S2A).

#### Enhanced syncytia formation and processing of BA.2.75 is determined by the N460K mutation

We further characterized the impact of BA.2.75-defining mutations on S fusogenicity and processing. Introduction of the N460K mutation into the BA.2 S drastically enhanced cell-cell fusion, with mean syncytia size 3.8-fold (p < 0.0001) higher than BA.2 (Fig. 4A-B; Fig. S2B). Conversely, introduction of the K460N reversion mutation into BA.2.75 significantly reduced cell-cell fusion, with mean syncytia size 4.3-fold (p < 0.0001) lower than BA.2.75 (Fig. 4C-D; Fig. S2C). We found that F157L and G257S in the BA.2 background, as well as the R152W reversion mutant in the BA.2.75 background, also exhibited modestly altered fusion activity (Fig. 4A-D). Importantly, the differences in membrane fusion between these mutants were not due to the surface expression level of S, as examined by flow cytometry (Fig. 4E-F; Fig. S2D-E). Consistent with enhanced fusion activity, introduction of the N460K mutation into the BA.2 S protein enhanced processing of S into the S1 and S2 subunits, as reflected in a S1/S ratio 40% higher than in BA.2 (Fig. 4G); a similar 70% increase in S2/S ratio was also observed (Fig. 4G). Conversely, introduction of a K460N reversion mutation into BA.2.75 reduced S protein processing by 20% (Fig. 4H). Thus, the N460K mutation in BA.2.75 enhances S processing, consistent with increased fusogenicity.

**Figure 4:**
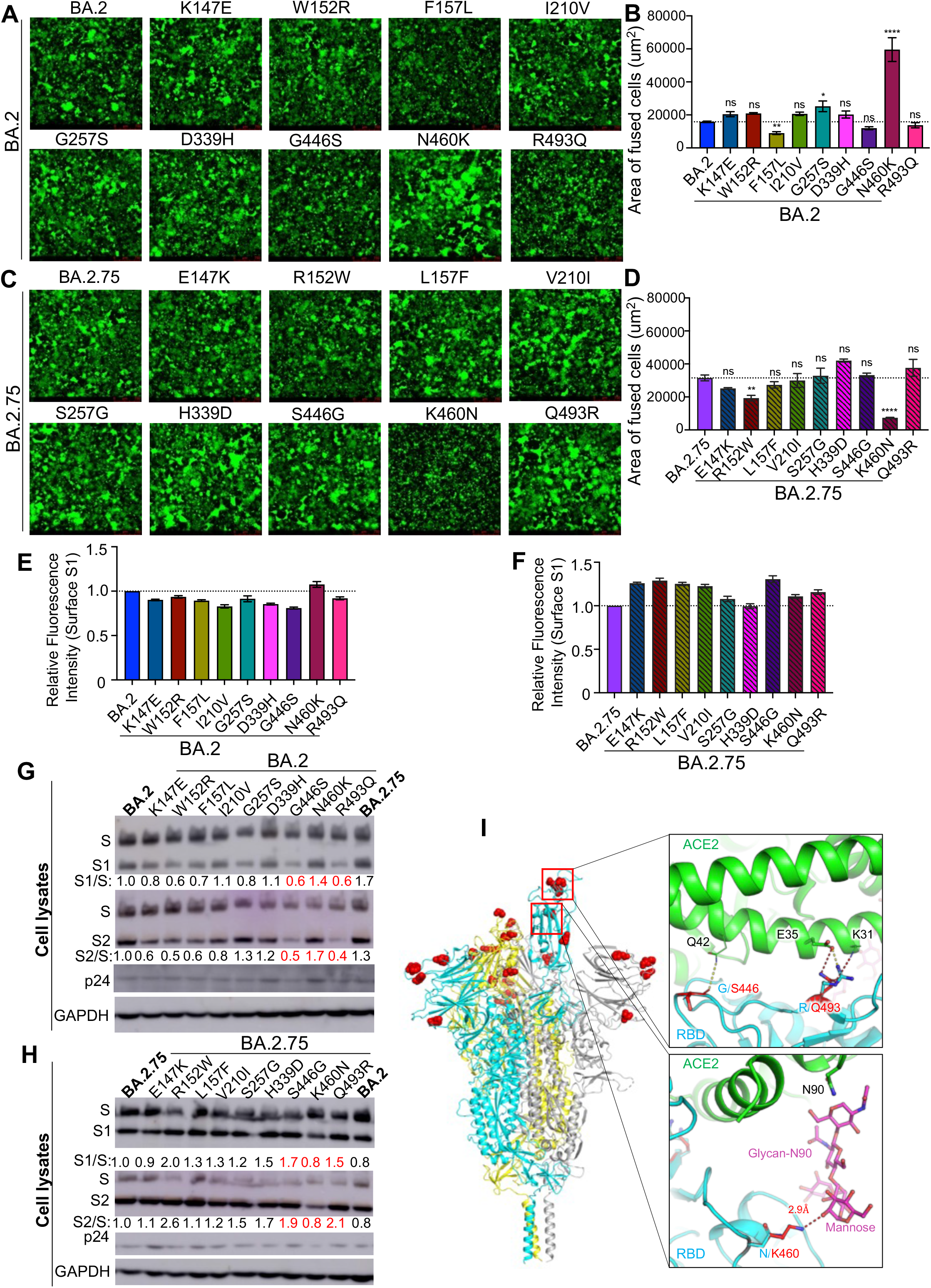
The N460K mutation determines enhanced cell-cell fusion and S processing of BA.2.75. (**A**) Fluorescence images displaying syncytia formation are presented for HEK293T-ACE2 cells 48 hr after co-transfection with a GFP expression construct and BA.2 single mutant S proteins. (**B**) Quantification of syncytia formation in panel (A) displays the mean syncytia size; bars represent means ± standard error, with significance relative to D614G determined by one-way ANOVA with Bonferroni multiplicity correction. (**C**) Fluorescence images displaying syncytia formation are presented for HEK293T-ACE2 cells 48-hrs after co-transfection with a GFP expression construct and BA.2.75 single reversion mutant S proteins. (**D**) Quantification of syncytia formation in panel (C) displays the mean syncytia size; bars represent means ± standard error, with significance relative to D614G determined by one-way ANOVA with Bonferroni multiplicity correction. (**E-F**) Quantification of relative S surface expression in transfected HEK293T cells for BA.2 single mutants (E) or BA.2.75 reversion mutants (F), as examined by flow cytometry; bars represent means ± standard error. (**G**) Pseudotyped lentivirus producer cell lysate was assessed for processing of S from BA.2 single mutants by probing with anti-S1 (T62), anti-S2, anti-HIV-1 p24, and anti-GAPDH. Band intensities were quantified in ImageJ and the ratios of S1/S and S2/S are displayed relative to the S1/S and S2/S ratios of BA.2. (**H**) Pseudotyped lentivirus producer cell lysate was assessed for processing of S from BA.2.75 reversion mutants by probing with anti-S1, anti-S2, anti-HIV-1 p24, and anti-GAPDH. Band intensities were quantified in ImageJ and the ratios of S1/S and S2/S are displayed relative to the S1/S and S2/S ratios of BA.2.75. (**I**) Structural modelling of Omicron BA.2.75 spike protein viewed as a ribbon. Mutations of BA.2.75 specific mutants are highlighted by red spheres. The RBD of the cyan spike protomer is in an “up” conformation. Upper inset: The mutation G446S reduces the backbone flexibility and possibly stabilizes the hydrogen bond between its carbonyl group and the residue Q42 on ACE2 receptor (green); the mutation R493Q abolishes the salt-bridge interaction with the E35 on ACE2 receptor and potentially forms two hydrogen bonds with E35 and K31. Lower inset: the mutation N460K enables formation of a hydrogen bond with the glycan-N90 on ACE2 receptor (green). In all cases, p-values are displayed as *p < 0.05, **p < 0.01, ****p < 0.0001, and ns for not significant.

### Structural modeling

To understand how BA.2.75 mutations contribute to functional changes, we created models of BA.2.75 spike protein and its complex with the ACE2 receptor using homology modeling (Fig. 4I). The G446S mutation does not appear to alter main chain interactions with the Q42 receptor residue; however, this mutation could reduce backbone flexibility, thus potentially stabilizing the specific interaction with ACE2, as well as spike integrity. The R493Q mutation would abolish a strong salt-bridge interaction with the E35 residue on the ACE2 receptor, which could reduce receptor binding affinity; however, this effect may be offset by the formation two new hydrogen bonds between the Q493 residue on spike and residues E35 and K31 on ACE2. Finally, N460K forms a new hydrogen bond with the glycan-N90 on ACE2 through an elongated side chain that reaches out to the alpha-1,3 mannose molecule on the N-linked glycan of the receptor residue N90, and this would likely enhance receptor binding affinity of BA.2.75.

## Discussion

The BA.2.75 subvariant is the latest in a series of Omicron variants to be identified. BA.2.75 has an alarming nine additional S mutations compared with BA.2, and preliminary reports suggest a slight growth advantage (Callaway, 2022; World Health Organization, 2022). These features portend that BA.2.75 could potentially overtake the BA.4/5 subvariants as the dominant circulating strain. Given this concern, it is critical to examine key features and novel phenotypes of BA.2.75, especially in its S protein. In this study, we show that BA.2.75 exhibits an increased neutralization resistance compared to ancestral BA.2, but has significantly lower neutralization resistance than BA.4/5 for 3-dose mRNA vaccinated HCWs as well as for hospitalized Omicron-wave patients. Critically, we demonstrate that the G446S and N460K mutations in the S protein of BA.2.75 underlie its enhanced neutralization resistance, while the R493Q mutation in BA.2.75, which is a reversion mutation, sensitizes it to neutralization. These findings could reflect the emergence of compensatory mutations to improve S function while maintaining neutralization resistance. Notably, the G446S mutation occurs in an epitope bound by class III neutralizing antibodies, rather than class II neutralizing antibodies that target the epitope of the R493Q mutation (Greaney et al., 2021). Structural analysis suggests that the side chain addition by G446S creates a steric clash with the CDR region of class III neutralizing antibodies, thus potentially hampering their recognition (Liu et al., 2022; Wang et al., 2022a). Hence, the exchange of these mutations may alter the susceptibility of BA.2.75 to class II and class III nAbs.

We further demonstrate that BA.2.75 exhibits enhanced S-mediated cell-cell fusion compared to BA.2, albeit to a lesser extent than BA.4/5. This enhanced triggering of BA.2.75 S-mediated fusion may reflect improved receptor utilization that is not present in earlier Omicron subvariants, consistent with several recent preprints (Cao et al., 2022; Saito et al., 2022; Wang et al., 2022a). Critically, we find that the N460K mutation present in BA.2.75 is essential for the enhanced fusion phenotype. This may relate to enhanced processing of N460K-containing S in virus producing cells, which would prime more cell surface-associated S for membrane fusion. While structural modeling did not provide an immediate explanation, the N460K mutation might enhance receptor utilization through a hydrogen bond with the receptor glycan N90. However, it is worth noting that this glycan interaction is mediated by a terminal mannose molecule, so it may not be easily observed in conditions of protein overexpression where glycosylation is often insufficient. G446S, on the other hand, may reduce the flexibility of loop 440-450, potentially enhancing overall spike thermostability, which likely decreases S processing efficiency. Furthermore, G446 is not well resolved in many apo spike structures, in line with its flexible local conformation. A more stable backbone loop conformation produced by the G446S mutation may reduce the energy cost for receptor engagement through hydrogen bond formation with Q42. Lastly, the loss of a strong salt-bridge interaction by the R493Q mutation is offset by the addition of two potential hydrogen bonds to the adjacent receptor residues, which could explain its modestly decreased fusion efficiency and processing. The contributions of these key residues to BA.2.75 replication kinetics in physiologically relevant human lung and airway epithelial cells needs to be carefully investigated. Further characterization of emerging SARS-CoV-2 variants will continue to aid our understanding of key features of SARS-CoV-2 evolution, spike biology, and immune evasion. Continued analysis of emerging variants also will improve ongoing public health responses and any potential reformulation of SARS-CoV-2 mRNA vaccine boosters.

Limitations of this study include a relatively small sample size for the boosted health care workers and the utilization of pseudotyped lentivirus for the neutralization assay rather than an authentic virus assay. However, our results for neutralization resistance are in accordance with several recent preprints (Cao et al., 2022; Gruell et al., 2022; Saito et al., 2022; Sheward et al., 2022; Wang et al., 2022b; Xie et al., 2022; Yamasoba et al., 2022a). Additionally, the lentiviral psedotype neutralization assay has been previously validated by assays with authentic SARS-CoV-2 (Zeng et al., 2020), and confirmed by numerous laboratories in the field. Future studies will focus on the biology and replication characteristics of BA.2.75 using variants isolated from human COVID patients.

## Author Contributions

S.-L.L. conceived and directed the project. P.Q. performed most of the experiments. J.P.E. assisted in experiments and contributed data processing and analyses. C.C. and R.J.G. provided clinical samples. P.Q., J.P.E., and S.-L.L. wrote the paper. K.X. performed homology modeling. Y.-M.Z, L.J.S., E.M.O. and K.X. provided insightful discussion and revision of the manuscript.

## Acknowledgements

We thank the NIH AIDS Reagent Program and BEI Resources for providing important reagents for this work and Xue Zou for assistance. We also thank the Clinical Research Center/Center for Clinical Research Management of The Ohio State University Wexner Medical Center and The Ohio State University College of Medicine in Columbus, Ohio, specifically Francesca Madiai, Dina McGowan, Breona Edwards, Evan Long, and Trina Wemlinger, for logistics, collection and processing of samples. In addition, we thank Sarah Karow, Madison So, Preston So, Daniela Farkas, and Finny Johns in the clinical trials team of The Ohio State University for sample collection and other supports.

## Funding

This work was supported by a fund provided by an anonymous private donor to OSU. S.-L.L., R.J.G., L.J.S. and E.M.O. were supported by the National Cancer Institute of the NIH under award no. U54CA260582. The content is solely the responsibility of the authors and does not necessarily represent the official views of the National Institutes of Health. J.P.E. was supported by Glenn Barber Fellowship from the Ohio State University College of Veterinary Medicine. R.J.G. was additionally supported by the Robert J. Anthony Fund for Cardiovascular Research and the JB Cardiovascular Research Fund, and L.J.S. was partially supported by NIH R01 HD095881. K.X. was supported by Path to K Grant through the Ohio State University Center for Clinical & Translational Science.

## Declaration of Interests

The authors declare no competing interests.

## Figure Legends

**Figure S1:**
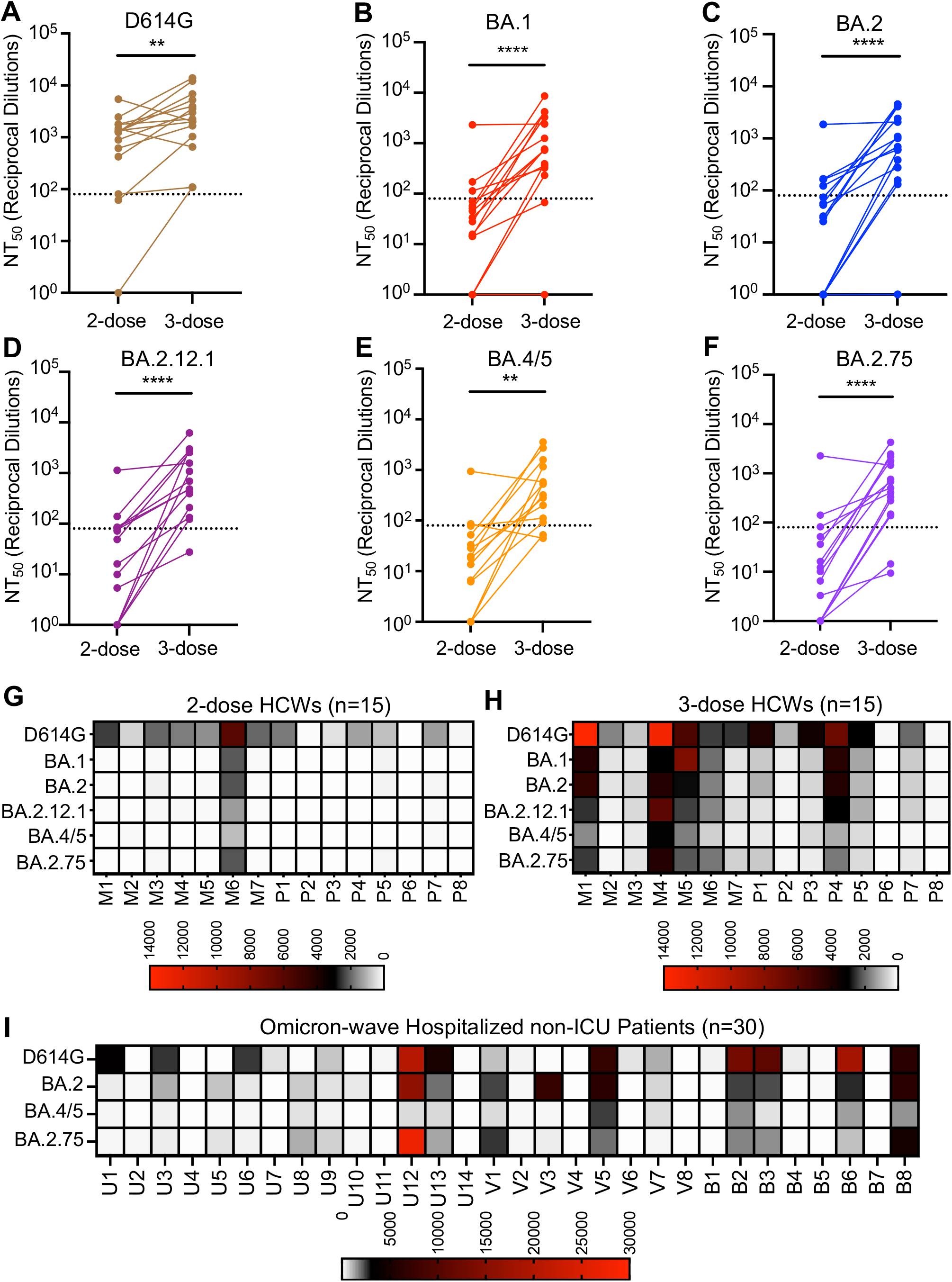
Neutralization of Omicron subvariants by vaccinee and COVID-19 patient sera, related to Figure 1. (**A-F**) Comparison of the neutralizing antibody titers in HCWs between 2-dose and 3-dose booster mRNA vaccination against the D614G (A), BA.1 (B), BA.2 (C), BA.2.12.1 (D), BA.4/5 (E), and BA.2.75 (F) variants. Lines connect samples from the same HCW, the dotted lines represent the limit of quantification (NT_50_ = 80), and significance was determined by paired, two-tailed Student’s t test with Welch’s correction. (**G-I**) Heatmaps display the nAb titers for HCWs 3-4 weeks after second mRNA vaccine dose (G), 1-12 weeks after mRNA vaccine booster dose (H), and for hospitalized Omicron wave COVID-19 patients (I). HCWs are indicated as ‘M’ for Moderna mRNA-1273 vaccinated or ‘P’ for Pfizer/BioNTech BNT162b2 vaccinated, and Omicron wave patients are indicated as ‘U’ for unvaccinated, ‘V’ for 2-dose vaccinated, and ‘B’ for vaccinated and boosted. P-values are represented as **p < 0.01 and ****p < 0.0001.

**Figure S2:**
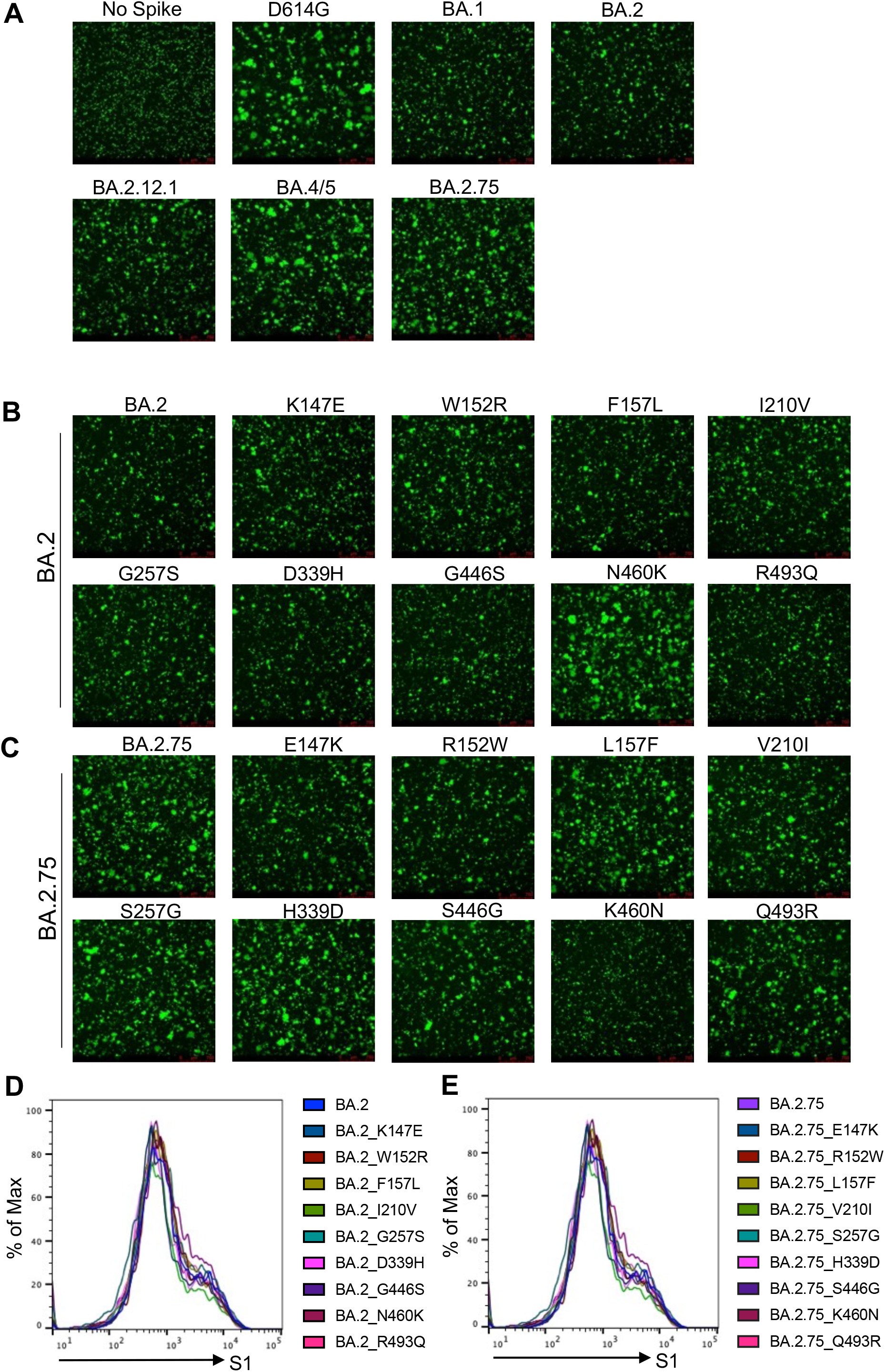
Syncytia formation and cell surface expression of Omicron subvariants, as well as BA.2- and BA.2.75-derived single mutants, related to Figures 3 and 4. (**A-C**) Fluorescence images displaying syncytia formation are presented for HEK293T-ACE2 cells 24 hr after co-transfection with a GFP expression construct and SARS-CoV-2 variant S proteins (A), BA.2 single mutants S proteins (B), or BA.2.75 single reversion mutant S proteins (C). (**D-E**) Histograms of surface staining with anti-S1 antibody of HEK293T cells expressing S proteins from BA.2 with single mutations from BA.2.75 lineage defining mutations (D) and from BA.2.75 with single reversion mutations from BA.2.75 lineage defining mutations (E).

## STAR Methods

### RESOURCE AVAILABILITY

#### Lead Contact

Further information and requests for resources and reagents should be directed to the lead contact, Dr. Shan-Lu Liu.

#### Materials Availability

Plasmids generated in this study are available upon request made to the lead contact.

#### Data and Code Availability

- NT_50_ values and de-identified patient information will be deposited to the National Cancer Institute SeroNet Coordinating Center. Additionally, NT_50_ values and de-identified patient information reported in this paper will be shared by the lead contact upon request.
- This paper does not report original code.
- Any additional information required to reanalyze the data reported in this paper is available from the lead contact upon request.

EXPERIMENTAL MODEL AND SUBJECT DETAILS

#### Patient Information

Sera were collected from the Ohio State University Wexner Medical Center health care workers (HCWs) under approved IRB protocols (2020H0228 and 2020H0527). Demographic information was self-reported and all subjects provided informed consent. Sera from 15 HCWs were collected 3-4 weeks after vaccination with a second dose of Moderna mRNA-1273 (n = 7) or Pfizer/BioNTech BNT162b2 (n = 8) vaccine, and 1-12 weeks after vaccination with a homologous booster dose. These HCWs ranged in age from 32 to 56 years (median 37 years) and included 6 female and 9 male HCWs. Analysis by age and gender could not be performed due to low sample number.

Sera were collected from patients 30 hospitalized for COVID-19 at the Ohio State University Wexner Medical Center under an approved IRB protocol (2020H0527). Sera were collected between early February and early March of 2022, during the Omicron wave in Ohio. Patients included 14 unvaccinated patients, 8 patients vaccinated with 2 doses of Moderna mRNA-1273 (n = 4) or Pfizer/BioNTech BNT162b2 (n = 4), and 8 patients vaccinated and boosted with Pfizer/BioNTech BNT162b2. This cohort included 11 female and 19 male patients. Patients ranged in age from 28 to 78 years (median 62 years).

#### Cell Lines and Maintenance

HEK293T (ATCC CRL-11268, RRID: CVCL_1926) and HEK293T-ACE2 (BEI NR-52511, RRID: CVCL_A7UK) cells were maintained in Dulbecco’s Modified Eagle’s Medium (DMEM) (Cibco, 11965-092) supplemented with 10% Fetal Bovine Serum (Signa, F1051) and 1% penicillin/streptomycin (HyCline, SV30010). Cells were maintained at 5% CO_2_ and 37°C.

#### METHOD DETAILS

##### Plasmids

Pseudotyped lentivirus was produced using a pNL4-3-inGluc lentivirus vector comprised of a ΔEnv HIV-1 backbone bearing a *Gaussia* luciferase reporter gene driven by a CMV promoter (Goerke et al., 2008; Zeng et al., 2020). SARS-CoV-2 S constructs bearing N- and C-terminal Flag tags were synthesized and cloned into a pcDNA3.1 vector by GenScript (Piscataway, NJ) by Kpn I and BamH I restriction enzyme cloning.

##### Pseudotyped lentivirus production and infectivity

Pseudotyped lentivirus was produced by transfecting HEK293T cells with pNL4-3-inGluc and S construct in a 2:1 ratio using polyethylenimine transfection. Pseudotyped lentivirus was collected at 48 hr and 72 hr after transfection. Collections were pooled and used to infect HEK293T-ACE2 cells to assess pseudotyped lentivirus infectivity. 48 hr and 72 hr after infection, infected cell culture media was assessed for *Gaussia* luciferase activity by combining 20 μL of media with 20 μL of *Gaussia* luciferase substrate (0.1 M Tris pH 7.4, 0.3 M sodium ascorbate, 10 μM coelenterazine). Luminescence was then immediately measured by a BioTek Cytation5 plate reader using BioTek Gen5 Microplate Reader and Imager Software (Winooski, VT).

##### Lentivirus neutralization assay

Pseudotyped lentivirus neutralization assays were performed as previously described (Zeng et al., 2020). Patient or HCW sera were 4-fold serially diluted in complete DMEM and pseudotyped lentivirus was added to neutralize for 1 hr (final dilutions: 1:80, 1:320, 1:1280, 1:5120, 1:20480, and no serum control). The pseudotyped lentivirus/sera mixtures were then transferred to HEK293T-ACE2 cells for infection. Then 48 hr and 72 hr after infection, infected cell media was assayed for *Gaussia* luciferase activity by combining 20 μL of cell culture media with 20 μL of *Gaussia* luciferase substrate. Luminescence was read immediately by a BioTek Cytation5 plate reader using BioTek Gen5 Microplate Reader and Imager Software (Winooski, VT). NT_50_ values were determined by least-squares-fit, non-linear regression in GraphPad Prism 9 (San Diego, CA).

##### Spike surface expression

HEK293T cells used to produce pseudotyped lentivirus were singularized by incubation in phosphate buffer saline (PBS) with 5 mM ethylenediaminetetraacetic acid (EDTA) at 37°C for 5 min and fixed 72 hr after transfection by incubation in 3.7% formaldehyde in PBS for 10 min. Cells were then stained with rabbit anti-S1 primary antibody (Sino Biological, 40150-T62) and anti-rabbit-IgG-FITC secondary antibody (Sigma, F9887). Samples were analyzed by a Life Technologies Attune NxT flow cytometer and data was processed using FlowJo v7.6.5 (Ashland, OR).

##### Syncytia formation

HEK293T-ACE2 cells were transfected with SARS-CoV-2 S constructs and a GFP expression construct. Cells were then imaged at 4x magnification 24 hr and 48 hr after transfection with a Leica DMi8 confocal microscope. Syncytia size was quantified using Leica Applications Suit X (Wetzlar, Germany) image analysis software. Three images were taken per sample with representative images being displayed.

##### Spike processing and incorporation

Pseudotyped lentivirus producing HEK293T cells were lysed by incubating in RIPA lysis buffer (50 mM Tris pH 7.5, 150 mM NaCl, 1mM EDTA, Nonidet P-40, 0.1% sodium dodecyl sulfate (SDS)) supplemented with protease inhibitor (Sigma, P8340) on ice for 30 min. Cell debris was pelleted and cell lysate was dissolved in 5x SDS-PAGE Laemmli buffer (312.5 mM Tris-HCl pH 6.8, 10% SDS, 25% glycerol, 0.5% Bromophenol blue, 10% β-mercaptoethanol). Pseudotyped lentivirus was purified by ultracentrifugation through a 20% sucrose cushion at 28,000 rpm and 4°C using a Beckman L-80 ultracentrifuge with TW-41 rotor. Pelleted pseudotyped lentivirus was resuspended in 1x SDS-PAGE Laemmli buffer. Cell lysate and purified virus were run on a 10% acrylamide SDS-PAGE gel and were transferred to a PVDF membrane. Membranes were blotted with anti-S1 (Sino Biological, 40150-T62), anti-S2 (Sino Biological, 40590-T62), anti-p24 (NIH ARP-1513), and anti-GAPDH (Santa Cruz Biotech, sc-47724) with anti-mouse-IgG-peroxidase (Sigma A5278) and anti-rabbit-IgG-HRP (Sigma, A9169) secondary antibodies. Blots were imaged with Immobilon Crescendo Western HRP substrate (Millipore, WBLUR0500) on a GE Amersham Imager 600. Band intensities were quantified using ImageJ (Bethesda, MD) image analysis software.

##### Homology modeling

Structural modeling of Omicron BA.2.75 spike protein and its complex with ACE2 receptor was conducted on SWISS-MODEL server with cryo-EM structure of SARS-CoV2 Omicron BA2 strain spike and complexes (PDB 7TNW and 7XB0) as templates. Glycan modeling, residue examination and rotamer adjustment were carried out manually with programs Coot (Cambridge, England) and PyMOL (New York, NY).

##### Quantification and statistical analysis

NT_50_ values were determined by least-squares-fit, non-linear-regression in GraphPad Prism 9 (San Diego, CA). NT_50_ values were log_10_ transformed for hypothesis testing to better approximate normality. Throughout, multiplicity was addressed by the use of Bonferroni corrections. Statistical analyses were performed using GraphPad Prism 9 (San Diego, CA) and are referenced in the figure legends and include one-way ANOVA (Fig. 3B and Fig. 4B and D), one-way repeated measures ANOVA (Fig. 1C-F, Fig. 2C-D), and a paired, two-tailed Student’s t test with Welch’s correction was used (Fig. S1A-F). Syncytia sizes were quantified by Leica Applications Suit X (Wetzlar, Germany). Band intensities (Figs. 3E and Fig. 4G-H) were quantified by ImageJ (Bethesda, MD) image analysis software.

## Notes

### Competing Interest Statement

The authors have declared no competing interest.

